# Wharton’s Jelly-derived mesenchymal stromal cells retain their immunophenotype and immunomodulating characteristics after transfection with polyethylenimine

**DOI:** 10.1101/2020.04.23.056663

**Authors:** Ana Isabel Ramos-Murillo, Elizabeth Rodríguez, Cristian Ricaurte, Karl Beltrán, Bernardo Camacho, Gustavo A. Salguero, Rubén Darío Godoy-Silva

## Abstract

**Background:** Wharton’s Jelly-derived mesenchymal stromal cells (WJ-MSCs) present several advantages over other sources of multipotent stem cells, not only because they are obtained from neonatal umbilical cord, which is considered a biological waste, but also display higher proliferation rate and low senescence at later passages compared to stromal cells obtained from other sources. In the field of tissue engineering, WJ-MSCs have a wide therapeutic potential, due to their multipotential capacity, which can be reinforced if cells are genetically modified to direct their differentiation towards a specific lineage; unfortunately, as primary cells, WJ-MSC are difficult to transfect. Therefore, the objective of the present work was to standardize a protocol for the transfection of WJ-MSCs using a cationic polymer. Such protocol is important for future developments that contemplate the genetic modification of WJ-MSCs for therapeutic purposes.

**Methods:** In this work, WJ-MSCs were genetically modified using polyethylenimine (PEI) and a lentiviral plasmid that encodes for green fluorescent protein (pGFP). To achieve WJ-MSCs transfection, complexes between PEI and pGFP, varying its composition (N/P ratio), were evaluated and characterized by size, zeta potential and cytotoxicity. At the N/P ratio condition where the highest transfection efficiencies were obtained, immunophenotype, immunomodulation properties and multipotential capacity of WJ-MSCs were evaluated.

**Results:** Here, we present the standardization of the transfection conditions of the WJ-MSCs in a monolayer culture with PEI. The concentrations of plasmid and PEI that have the best transfection efficiencies were established

**Conclusions:** Transfection with PEI doesn’t affect immunophenotype, immunomodulatory properties and differentiation capacity of WJ-MSCs.

## Background

Human mesenchymal stromal cells (hMSCs) are used in a wide variety of therapeutic applications due to its regenerative capacity and multipotentiality [1–3]. Tissue engineering is one of the most promising fields of application due to hMSCs capacity to differentiate in several cell lineages, such as adipocytes (fat tissue), osteocytes (bone) and chondrocytes (cartilage)[4]. This multipotentiality is mediated by several inductive factors which are necessary to reach the differentiation to a specific lineage. hMSCs differentiation can be reached through the addition of inductive factors in the medium. In 1998, Johnstone *et al* developed a methodology to reach *in vitro* chondrogenesis of bone marrow-derived mesenchymal progenitor cells that includes the use of high glucose medium, a premix of ITS (insulin, transferrin, and selenous acid), ascorbate 2-phosphate and Transforming Growth Factor-β (TGF-β) 1 or 3 [5]. In 1999, Pittenger *et al* demonstrated the multilineage potential of adult hMSCs and induced adipogenic differentiation by treatment with 1-methyl-3-isobutylxanthine, dexamethasone, insulin, and indomethacin. To promote osteogenesis, hMSCs were treated with dexamethasone, b-glycerol phosphate, and ascorbate and in the presence of 10% v/v Fetal Bovine Serum (FBS) [6].

From then until now, several authors have developed strategies not only to differentiate stem cells but also to find new sources of obtaining them. In 2004, Wang *et al* evaluated 30 fresh human umbilical cords obtained after birth and demonstrated that mesenchymal cells from this tissue (UC-hMSCs) upregulated expression of adhesion-related molecules (CD105, CD44) and integrin markers (CD51, CD29); however, such was not the case for markers of hematopoietic differentiation (CD45, CD34). Additionally, the authors successfully differentiated those cells to the three lineages (osteoblasts, adipocytes and chondroblasts) under suitable culture conditions. Those results suggest that stroma cells from Wharton’s jelly are similar to mesenchymal stem cells (MSCs) [7] and can be used as a source of stem cells for tissue engineering. WJ-MSCs have some advantages over bone marrow cells, considering that they can be stored and do not require perfect human leukocyte antigen (HLA) tissue matching and they are permissive to allogeneic transplantation [8].

Several investigators have found that the regenerative capacity of therapies based on hMSCs is deficient when cells are employed without the proper growing factors, due to a poor survival and lack of differentiation. To overcome those difficulties, hMSCs can be modified through genetic engineering to improve their survival rate and increase the secretion of differentiation factors to induce their differentiation to a specific lineage.

Gene therapy consists of the introduction of specific genes into patient cells to produce a therapeutic effect [9]. This modification can be carried out mainly through two strategies: (1) using viral vectors as lentivirus, adeno-associated virus or adenovirus among others, and (2) using non-viral vectors. Viral modification allows efficient transfer of the transgene and if in the case of retrovirus, constitutive production of the protein of interest; however, its methodology might increase the risk of insertional mutagenesis and/or the activation of oncogenes. The use of viral vectors is widely used for clinical applications but is subjected to rigorous controls, which implies high production costs [10].

On the other hand, non-viral vectors are preferred when transient modifications are expected. Liposomes and cationic polymers are widely used as they are easily scalable and are suitable for their use in several modification approaches. Nevertheless, the major disadvantage of non-viral vectors is the low efficiency of transfection [11]. While with viral vectors it is possible to reach efficiencies above 90% in hMSCs, in the case of non-viral vectors its value is reduced to 10-15%.

In 1995, Boussif and collaborators proposed PEI as a potential transfection agent, considering that polycations with buffer capacity at physiological pH are effective transfection agents *per se*, that is, without adding targeting or membrane rupture agents [12]. Since then, more than 2000 articles have been published related to the use of PEI in transfection assays or as a gene-releasing agent (supply of genes) and in the previous years the use of PEI has been explored as a transfection agent of mesenchymal stromal cells, with more than 135 publications in this topic involving mesenchymal stromal cells. Despite this, there are few studies that explore the use of branched PEI 25 kD as a transfection agent of mesenchymal cells from Wharton’s Jelly (WJ-MSCs) [13, 14].

According to the ISCT [15, 16], WJ-MSCs can be classified under the name of stromal cells, given that the potential capacity for multilineage differentiation (adipogenesis, chondrogenesis and osteogenesis) [17, 18], among others. Wharton’s Jelly-derived mesenchymal stromal cells (WJ-MSCs) present several advantages over other sources of multipotent stem cells, not only because they are obtained from neonatal umbilical cord, which is considered a biological waste, but also display higher proliferation rate and low senescence at later passages compared to stromal cells obtained from other sources. In the field of tissue engineering, WJ-MSCs have a wide therapeutic potential, due to their multipotential capacity, which can be reinforced if cells are genetically modified to direct their differentiation towards a specific lineage; unfortunately, as primary cells, WJ-MSC are difficult to transfect. Therefore, the objective of the present work was to standardize a protocol for the transfection of WJ-MSCs using PEI and the to evaluate if transfection with PEI affect those important characteristics of stromal cells such as: immunomodulatory properties, differentiation capacity and expression of specific markers.

## Materials and methods

### Materials

#### Cell culture

Growth media Dulbecco’s Modified Eagles Medium -DMEM (Gibco, Life Technologies Corp, USA), and Fetal Bovine Serum FBS (Biowest, USA and Sigma Aldrich, USA) were employed for HEK-293 and WJ-MSCs culture. RPMI (Gibco, Life Technologies Corp, USA) was used for the culture of peripheral blood mononuclear cells (PBMCs). Dimethyl sulfoxide (DMSO) from Sigma Aldrich (USA) was used at 5%v/v for cell freezing.

#### Transfection

Branched polyethylenimine (PEI)(25 kDa) from Sigma (USA) was employed in all transfection assays. Nucleus staining: DAPI (4′,6-diamidino-2-phenylindole) was used at 300nM (D9542, Sigma Aldrich, USA) in a Phosphate Buffered Solution (PBS) at pH 7.4.

#### Cell viability

440 µM Resazurin sodium salt (121519, PanReac AppliChem, Darmstadt, Germany) in PBS at pH 7.4 was used to determine the metabolic activity of cells as a measurement of cell viability.

#### *E. coli* DH5-*α* culture

Luria Bertani (LB) medium (L3152) and ampicillin sodium salt (A0166) were acquired from Sigma Aldrich (San Luis MO, USA) and agar-agar (A0949) was supplied by PanReac AppliChem (Darmstadt, Germany).

### Expansion of WJ-MSCs and HEK

WJ-MSCs were obtained from umbilical cord donors (number 40 and 148) from the Advanced Therapies Unit at the Instituto Distrital de Ciencia Biotecnología e Innovación en Salud following informed consent by the mothers. WJ-MSCs were cultured in DMEM supplemented with 10% FBS. Cells from cord number 40 were passaged at 80% confluency and expanded to passage 9, and then were cryopreserved (3 x10^6^cells/vial) (2 ml DMEM, 10% SFB and 10% DMSO) in liquid nitrogen. Cells from cord number 148 were passaged at 80% confluency and expanded to passage 4 and then frozen as previously described. Human embryonic kidney cells 293 (HEK-293) were cultured in DMEM-F12 supplemented with 10% FBS. Cells were passaged at 80% confluency and expanded to passage 12, and then frozen to posterior use.

### Plasmid propagation

*E. coli* DH5-α cells kindly donated by Dr. Velasquez laboratory (National University of Colombia) were employed. Cells were cultured in 100 ml shake flasks containing 50 ml of Luria-Bertani medium (Tryptone, NaCl and yeast extract) at 37°C on an orbital shaker at 230 rpm and growth was halted when the optical density at 600 nm (OD600) was approximately 1.2. To obtain competent bacteria, *E. coli* cells were recovered by centrifugation at 3500*g* and were resuspended twice in 10 ml of a calcium solution at 0°C (CaCl_2_ 60 mM, Glycerol 15%, Pipes 10 mM, pH: 7.0). Then, cells were resuspended in 2 ml of the same calcium solution and aliquoted (80 µL) and stored at −20°C. The competent *E. coli* cells were employed to obtain transformed *E. coli* cells with a plasmid (RRL.SIN.cPPT-hCMV-eGFP) coding for GFP (pGFP) and resistance to ampicillin. Briefly, competent cells were mixed with 5 µL of pGFP (1 µg) and were placed in an ice bath for 10 minutes, then in a 42°C bath for 2 minutes and finally 10 more minutes in an ice bath. Thereupon, competent cells were diluted with 500 µL of LB with ampicillin (100 µg/µL) and were seeded in solid media (LB media (25g/L), agar-agar (12g/L) and ampicillin (100 µg/µL)) using a Drigalski spatula. Competent cells were incubated at 37°C for 24-48 hours until visible colonies were visibly formed by transformed *E. coli* cells.

*E. coli* cells from colonies were seeded in LB media with ampicillin and were incubated at 37°C and 230 rpm by 16 hours. Then 1 ml of this culture was used to inoculate 150 ml of LB media with ampicillin. Transfected cells were cultured until an optic density at 600 nm (OD600nm) of 1,2 was reached. At this moment, *E. coli* cells were recovered by centrifugation and pGFP was purified using ZymoPURE™ Plasmid Maxiprep Kit (catalog #D4203, Zymo Research, Irvine, USA). The DNA concentration was measured in a NanoDrop™ 2000/2000c Spectrophotometer (ThermoScientific®, USA).

### PEI and pGFP nanoparticle formulation and physicochemical characterization

Transfection efficiency was evaluated at different N/P ratios (molar ratios of nitrogen and phosphorus in PEI and DNA, respectively). Grayson et al (2006) and Ahn et al (2008) calculate N/P ratio assuming that 43.1 g/mol (molecular weight of each repeating unit of PEI: C2H5N), containing one nitrogen atom and 330 g/mol (molecular weight average of each repeating unit of DNA: each nucleotide), containing one phosphorus atom [19, 20]. In this work, N/P ratio was calculated considering only primary amine nitrogen from branched 25kDa PEI, considering that 1 mol of PEI holds 0.25 mol of primary amine nitrogen [21] and using overall nitrogen content measurements in a Total Organic Carbon/Total Nitrogen (TOC/TN) analyzer (AnalytikJena, Jena, Germany).

Different N/P ratio were evaluated using 400 ng of pGFP. Detailed calculations of N/P ratio are presented in table S1 (Supplementary Material). pGFP and PEI concentrations were adjusted to mix always the same volume of each solution. PEI was added to DNA and the solution was vortexed immediately. Then, the solution was incubated at 37° for 30 minutes to favor complex formation. Nanoparticle size was measured through Dynamic Light Scattering (DLS) and zeta potential by Laser Doppler Micro-electrophoresis to evaluate the effect of N/P ratio on size (Zetasizer Nano ZS from Malvern Panalyticial (UK)). Complex formation was also evaluated by electrophoresis in agarose gel. Briefly, 0.79 g of Ultrapure^TM^ agarose (ThermoScientific) in 100 ml of Tris-acetate-EDTA (TAE) buffer was used. A Biorad power supply was employed to apply 100V through the agarose gel for 30 minutes. A 1kb marker was used as molecular weight marker and BlueJuice^TM^ (ThermoFisher Scientific, USA) was used as loading buffer. Nanoparticle size and zeta potential assays were carried out by triplicate. Distilled water was filtered through a 0.2 µm membrane and used as solvent.

### Measurement of metabolic activity

Cell viability was determined by measuring the metabolic activity, through the reduction of the sodium salt of resazurin (C12H6NNaO4) which is a key indicator of reduction-oxidation (redox) reactions. Because of the metabolic activity of the viable cells, resazurin is reduced to resorufin resulting in a color change from blue to highly fluorescent fuchsia in response to a chemical reduction of the culture medium. The reduction of resazurin can be monitored by measuring fluorescence or absorbance [22]. The mechanism of reduction of resazurin through viable cells is based on the use of NADH and NADPH as electron sources. The reaction occurs due to the action of mitochondrial, microsomal enzymes or enzymes of the respiratory chain [23]. In short, cells were cultured in a 24 well plate and the culture medium was removed and a resazurin solution (10% v/v resazurin 440 μM in supplemented media) was added to each well after cells adhesion. The intensity of the fluorescence was measured hourly at an excitation wavelength of 530 nm and emission of 590 nm. The percentage of reduction of resazurin was calculated as a relation between the Relative Fluorescence Units (RFU) of the experiment and their respective controls (equation 1):

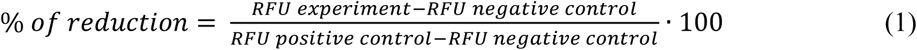

Resazurin diluted in supplemented medium was completely reduced to resorufin by autoclaving at 121°C and it was used as positive control. Same dilution without autoclaving was used as negative control.

### Assessment of polyethylenimine (PEI) cytotoxicity on transfected WJ-MSCs and HEK-293

Branched PEI (25 kDa) was employed as transfection agent. A PEI stock at 10 µg/µl was prepared in distilled water (DW), and pH was adjusted at 3.15 with chloride acid (HCl). PEI stock solution was filtered using a 0.2 µm filter. PEI stock at 10 µg/µl was diluted in DW to obtain a working solution of PEI at 1µg/µL. PEI working solution was aliquoted at 500 µl and frozen at −20°C. For all experiments, fresh working PEI solution was used to avoid freezing and thawing several times the same stock. PEI cytotoxicity was evaluated through resazurin assay. For this purpose, cells were seeded in a 48 well plate at a density of 4.5×10^4^ cells per well for WJ-MSCs and 1×10^6^ cells per well for HEK, 24 hours prior to cytotoxicity assay. PEI dilutions were prepared in DW and then 10 µl of PEI solution were diluted in 90 µl of non-supplemented DMEM F12. Cells were treated with different concentration of PEI for 4 hours at 37°C and 5% of CO_2_, then media with PEI was removed, and 100 µl of resazurin solution was added and fluorescence was quantified each hour at the excitation and emission wavelengths of 560 and 590 nm respectively, using a Cytation 3 microplate reader (Biotek®, Vermont,USA). All assays were conducted in triplicate. Cells were treated with non-supplemented media to evaluate the serum deprivation effect.

### Growth kinetics of WJ-MSCs

Growth of WJ-MSCs was followed using two different initial cell densities. For the first one, WJ-MSCs at passage 6 were seeded at 2000 cells/cm^2^ and for the second one 9000 cells/cm^2^ were cultured in a 48-well plate. After 24 hours, medium was removed from 6 wells and 100 µL of resazurin solution was added to each one to follow the metabolic activity through the change in fluorescence for 12 hours. Well plates were always protected from light and fluorescence measurements were carried out at 37°C and 5% CO_2_ at Cytation 3 (Cell Imaging Multi-Mode Reader) from Biotek® (USA). Next day, same procedure was followed using six different wells. This protocol was repeated until day 6.

### *In vitro* evaluation of transfection efficiency of PEI–pGFP complexes in WJ-MSCs

WJ-MSCs at passage 6 were plated at a seeding density of 1×10^4^ cells/cm^2^ and HEK-293 at a density of 1×10^5^ cells/cm^2^ in 24-well adherent plates (SPL, Korea) 48 h prior to transfection. Complexes prepared as previously described were incubated for 30 minutes and before the transfection, media was removed from wells and complexes formed by PEI and pGFP was added to cells and the plate was centrifuged at 280*g* for 5 minutes. Cells were incubated for 4 hours with the complexes and then media was removed and DMEM-F12 supplemented was added. Transfection efficiency was evaluated trough fluorescence using a Cytation 3 microplate reader (Biotek®, Vermont, USA). Briefly, transfected cells were trypsinized and resuspended in serum free media. On the one hand, cells were stained with Trypan blue exclusion technique and total HEK cells per well and viability was measured. On the other hand, transfected cells were resuspended in 100 μl of serum free media and fluorescence intensity (Relative Fluorescence Units (RFU)) was measured and normalized with respect to total HEK cells.

Different N/P ratios were evaluated to analyze their effect on WJ-MSCs transfection efficiency. After a total incubation time of 48 h, cell transfection was visualized by fluorescence. Then, WJ-MSCs were trypsinized and harvested by centrifugation. Cells were resuspended in 100 μl of PBS 1X and evaluated by flow cytometry. Data were analyzed with Kaluza Analysis 2.0 (Beckman Coulter, USA) software. Untreated cells and cells treated with naked, non-complexed plasmid were employed as controls. The data is represented as the mean of the standard deviation (SD) for each treatment carried out in triplicate.

### Immunophenotype of WJ-MSCs transfected with PEI

In order to evaluate the presence of characteristic cell surface markers of hMSCs and the absence of those of hematopoietic cells, WJ-MSCs after 48 hours of transfection with PEI at N/P ratio of 3.5 (720 ng PEI/90.000 cells) were recovered by trypsinization. The expression of MSC-related cell surface antigens was assessed by flow cytometry using the membrane markers CD90 (APC), CD73 (PECy7), CD105 (PE), CD45 (APC/Cy7), CD34 (PerCP-Cy5.5), HLA-DR (Pacific Blue) y HLA-ABC (FITC). Cells were incubated for 30 min at 4°C, centrifuged at 300g for 6 min and resuspended in 0.2 mL PBS. The procedure was performed in a flow cytometer FACSCanto II (Becton Dickinson, San Jose, USA) and data was analyzed with FlowJo vX.7.0 data analysis software package (TreeStar, USA).

### Immunomodulation properties of WJ-MSCs transfected with PEI

To assess the ability of transfected WJ-MSCs to inhibit the proliferation of CD3+ T lymphocyte an immunomodulation assay was carried out. Peripheral blood mononuclear cells (PBMCs) were cultured in RPMI media supplemented with FBS at 10% v/v and were used as a source of T lymphocyte cells. WJ-MSCs were transfected following the procedure describe previously (N/P ratio of 3.5). After 48 hours of transfection, cells were trypsinized and seeded into 24-well plates at 5×10^4^ cells per well and cultured for 24 hours. Then 5×10^5^ PBMCs activated (A) with CD2, CD3 and CD28 monoclonal antibodies (beads T cell activation/Expansion Kit, Miltenyi Biotec GmbH, Bergisch Gladbach, Germany) were co-cultured with WJ-MSCs (A/ WJ hMSCs) and incubated at 37ºC with 5% humidified CO_2_ for 5 days. PBMCs activated (A) and without activation (NA) were used as control. After 5 days of culture, cells were recovered from wells and were incubated at 4°C for 30 minutes with antiCD3 antibody (Biolegend, CA, USA) and then were examined by flow cytometry.

### Multipotential capacity of WJ-MSCs

The multipotential capacity of WJ-MSCs after transfection with PEI at N/P ratio of 3.5 (720 ng PEI/90.000 cells) was examined by cultivating transfected cells in secondary cultures using osteogenic, chondrogenic, and adipogenic differentiation media. To induce adipogenic differentiation, transfected cells were seeded into 24-well plates at 5×10^4^ cells per well and cultured until 60% confluency was achieved, at which point they were incubated in adipogenic induction medium (Stromal Pro Adipogenesis Differentiation Kit, Life Technologies, USA). Media was changed every 3 days and after 14 days, cells were fixed in 4% paraformaldehyde (Sigma Aldrich, St Louis, MO, USA) prior to staining of lipid vacuoles with Oil Red O (Sigma Aldrich, St Louis, MO, USA) staining. For osteogenic differentiation, transfected cells were similarly seeded, as previously described in osteogenic differentiation medium (Stromal Pro Osteogenesis Differentiation Kit, Life Technologies). Calcium deposition was analyzed using alizarin red-S (Sigma Aldrich, St Louis, MO, USA) staining. For all controls (non-differentiation conditions), cells were cultured in DMEM culture medium. In all cases, observation was made microscopic and photographic record.

### Statistical analysis

Statistical analyses were performed using GraphPad Prism XVII software. One-way ANOVA was used for analysis of variance with Tukey’s post hoc test to compare between groups. Numerical and graphical results are displayed as mean ± standard deviation. Significance was accepted at a level of p < 0.05.

## Results

In the present study, we evaluated the non-viral gene modification of human umbilical cord Wharton’s Jelly-derived mesenchymal stromal cells (WJ-MSCs) using polyethylenimine (PEI), a cationic polymer widely used in transfection assays. Different PEI/pGFP complexes were prepared by electrostatic interactions between the negatively charged phosphates on the DNA (pGFP) and the free amino groups on PEI. The formed complexes were characterized and evaluated in HEK cells and umbilical cord mesenchymal stromal cells (WJ-MSCs) for transfection. The effect of transfection with PEI in the multipotential differentiation capacity and immunomodulatory properties of WJ-MSCs was also studied.

### pGFP and PEI/pGFP complex characterization

Molecular weight of pGFP was obtained by electrophoresis. According to the electrophoretic gel showed in Fig. 1A, GFP plasmid size was approximately 8kb. Furthermore, it is observed that when the plasmid is not digested, several bands can be distinguished, which correspond to the whole plasmid but do not allow to establish its molecular weight.

**Fig. 1.**
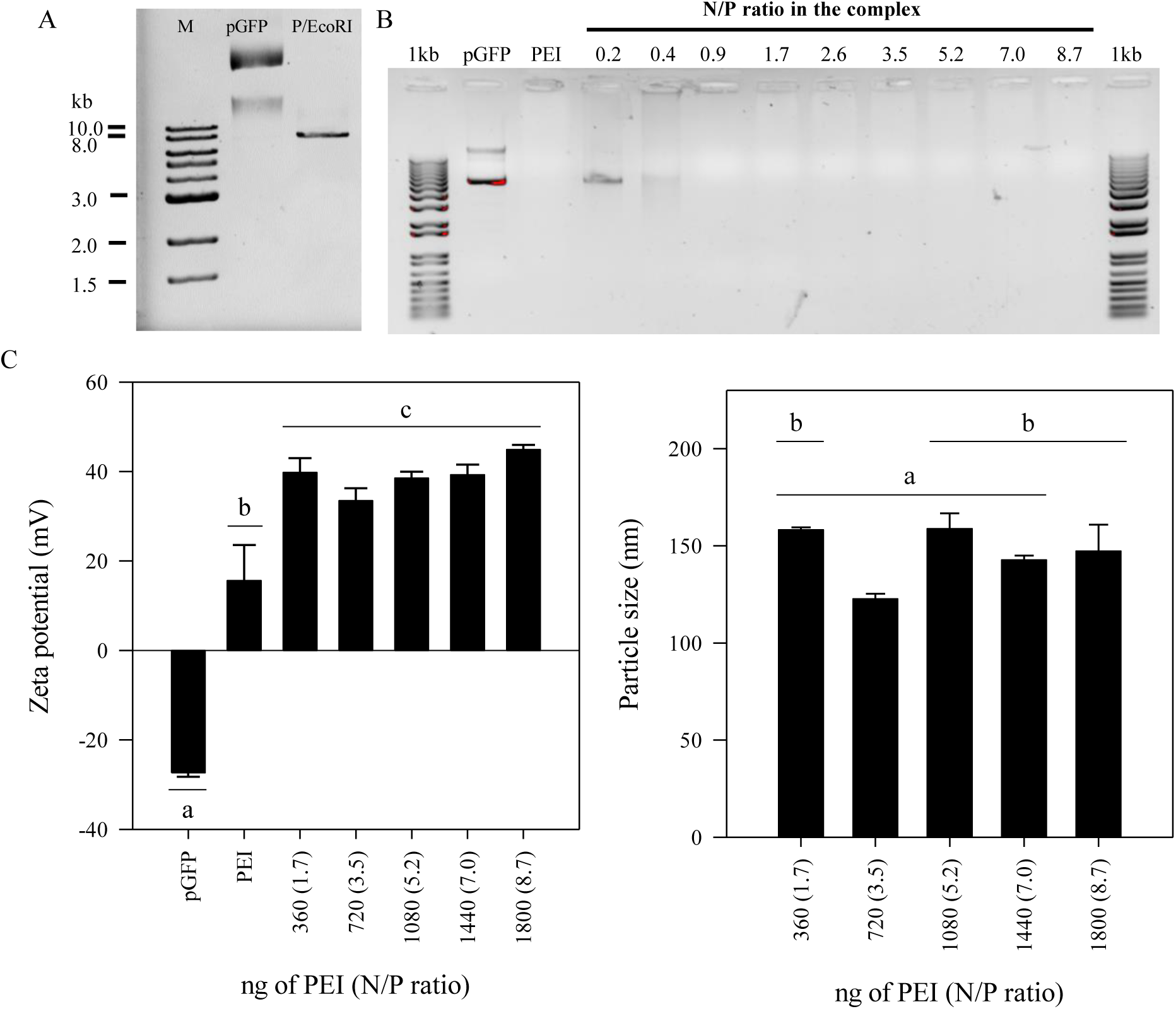
Evaluation of PEI/pGFP complexes formation. 400 ng of pGFP plasmid were mixed with PEI solutions in DW at different concentrations, to obtain complexes at several N/P ratios. N/P=0.9 correspond to 180 ng of PEI (1.04 moles of nitrogen from primary amines). (A) Gel retardation assay of pGFP to establish its molecular weight, (B) PEI/pGFP complexes formation by gel retardation assay, (C) Effect of N/P ratio on Zeta potential and (D) complexes size. (^a,b,c^) denotes significance (n = 3, p < 0.05) in comparison to all groups in the same time point.

To evaluate the formation of complexes of PEI and pGFP, an electrophoresis analysis was carried out. Fig. 1B shows the electrophoretic patterns of complexes prepared at different N/P ratios. Total content of nitrogen in the sample was measured by high-temperature combustion (multi N/C 3000, AnalytikJena, Germany). Total nitrogen content in the PEI solution (50mg/l) was 14.60 mg/l ±1.16%.

According to the content of primary amines in PEI, different N/P ratio were evaluated (0.2, 0.4, 0.9, 1.7, 2.6, 3.5, 5.2, 7.0 and 8.7). Migration of plasmid DNA (pGFP) was delayed by complexation with PEI. When N/P ratio was equal to or lower than 0.9, some bands were displaced through the agarose gel, similarly to the plasmid alone, indicating in these conditions that PEI concentration is not enough to complex the plasmid. These results could suggest than N/P ratios lower than 0.9 are not appropriate for transfection assays. Zeta potential and particle size of PEI/pGFP complexes were evaluated by Laser Doppler Micro-electrophoresis and Dynamic Light Scattering (DLS) (Fig. 1C and Fig. 1D) respectively. In zeta potential measurements, an electric field is applied to a solution of molecules or a dispersion of particles, which then move with a velocity proportional to their zeta potential. This velocity is measured using a laser interferometric, which enables the calculation of electrophoretic mobility, and thus, the zeta potential and its distribution. DNA plasmids are molecules with a highly negative surface charge due to the presence of phosphate groups. Measured zeta potential of the pGFP plasmid used in this work was approximately −27±0.9 mV in a plasmid solution of 400ng pGFP/ml in distilled water (DW). Branched PEI is a molecule with free amino groups that give it a positive net charge. 1.2 ml of a solution of Branched PEI at 33.3 µg/ml in DW was also evaluated, and zeta potential measurements oscillated between 6 and 26mV, finding a wide variation among the results. When the interaction between PEI and pGFP starts to form complexes, PEI concentrations lower than 180ng (N/P ratio of 0.9) are unable to retain DNA and no particles are formed. When complexes are formed at N/P ratio higher than or equal to 1.7, the size and the zeta potential of the complexes remain constant between 130-180 nm and 36-46 mV, respectively.

### Cytotoxicity of pGFP and PEI/pGFP complexes

The effect of PEI on the metabolic activity of HEK cells and WJ-MSCs was evaluated by resazurin assay technique. Cells were treated with different PEI solutions in culture media without supplementation. It was found that HEK cells are more resistant to PEI toxicity than WJ-MSCs. PEI concentrations higher than 1800 ng/100.000 HEK cells (180ng/10000 HEK cells) reduce cell metabolic activity between 20-30% as compared to WJ-MSCs where the same concentration reduces cell viability (Fig. 2A and Fig. 2B). It was observed that concentrations between 540 ng and 1080ng/45,000 WJ-MSCs (120 ng/10000 WJ-MSCs) reduce the metabolic activity around 20-40%.

**Fig. 2.**
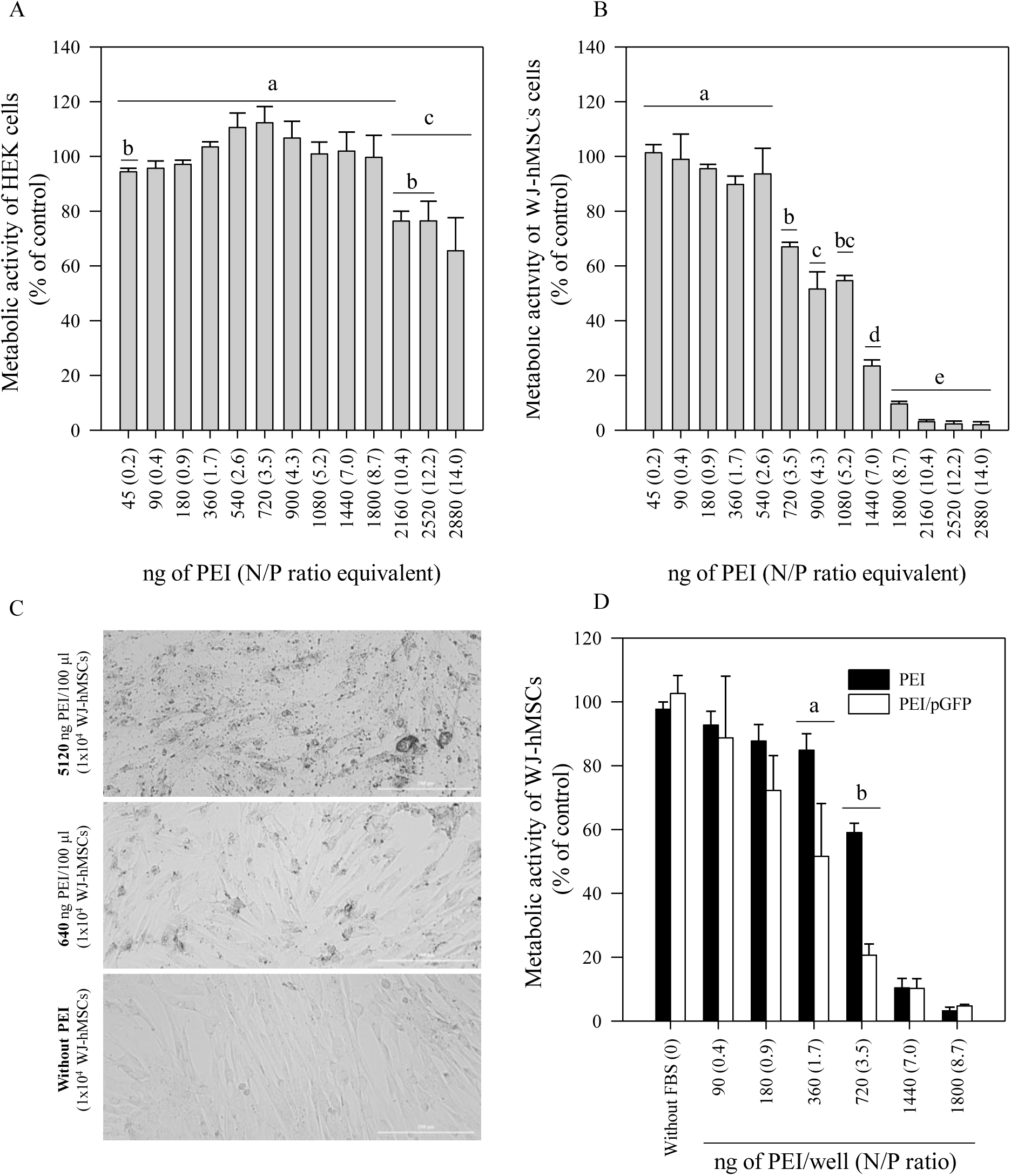
Effect of PEI concentration at metabolic activity of cells in media without supplement, after 4 hours of treatment with different PEI concentrations. (A) 1×10^5^ HEK cells/100 µL, (B) 4.5×10^4^ WJ-MSCs/100 µL), (C) WJ-MSCs non-treated (without PEI) and treated with 640 and 5120 ng of PEI/well after 24 hours and (D) 4.5×10^4^ WJ-MSCs /200 µL treated with complexes formed with 800 ng of pGFP. (^a,b,c,d,e^) denotes significative differences (n = 3, p < 0.05) in comparison to all groups in the same time point.

Above this concentration, the metabolic activity is reduced approximately 80% and concentrations equal to or higher than 1800 ng lead to cell death. Bright field photographs of non-treated WJ-MSCs and those treated with 640 and 5120 ng of PEI/well after 24 hours are presented in Fig. 2C. Cytotoxicity of PEI/pGFP complexes was evaluated (Fig. 2D). Significant differences were found between PEI alone and PEI complexed with pGFP at the same concentration. At 1.7 and 3.5 N/P ratios, complexes were more cytotoxic than non-complexed PEI. These results can be associated with the cellular internalization of complexes.

### Transfection assay with PEI and pGFP complexes

Transfection with PEI was evaluated in HEK cell line and WJ-MSCs. HEK cell line reached confluence at a concentration of approximately 3×10^6^ cells/cm^2^, while WJ-MSCs did at 4.5×10^5^cells/cm^2^. Different PEI concentrations were used for transfection assays, keeping the concentration of pGFP constant.

### HEK transfection

HEK cells are widely used as control in transfection experiments due to their transfectability. The effect of PEI concentration in transfection efficiency was evaluated (Fig. 3). Different PEI/pGFP mass ratios were evaluated to establish the best combination of PEI and pGFP with less toxicity. An increase in PEI concentration did not reduce cell viability but reduced cell concentration per well, as shown in Fig. 3A. These results suggest a cytotoxic effect, reflected in cell concentration reduction when N/P ratio is equal to 9.4.

**Fig. 3.**
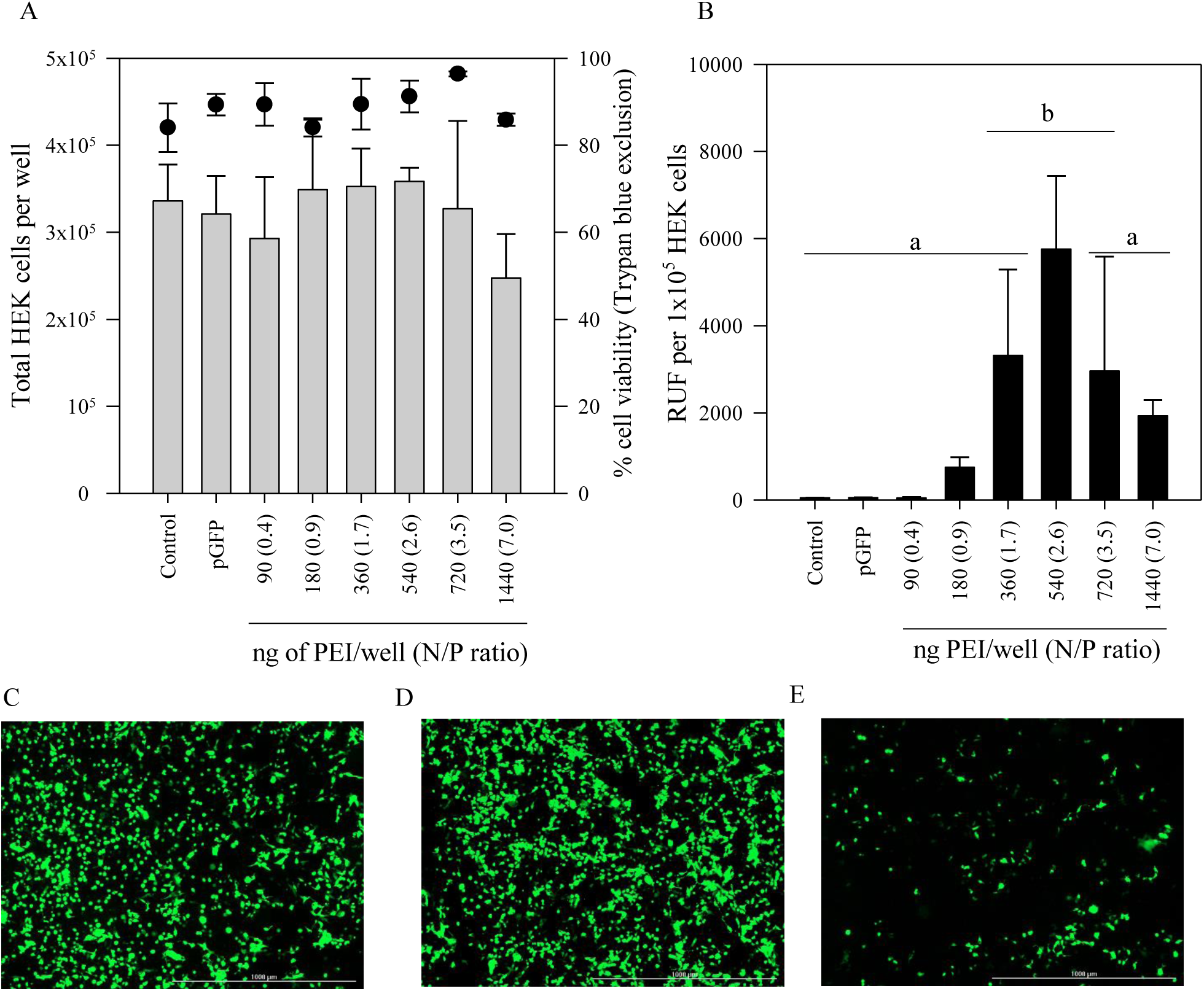
Effect of PEI concentration in HEK cells transfection. (A)Total HEK cells (left -*y* axis) and percentage of viable cells (right -*y* axis) by Trypan blue exclusion after 4 hours with PEI/pGFP complexes treatment. (B) Relative units of fluorescence (RFU) of HEK cells after 48 hours of transfection. (C, D, and E) Fluorescent microscope images of the GFP expressing HEK 48 h post transfection. N/P ratios of 1.7, 2.6 and 3.5 respectively. Scale bar, 1000 um. (^a,b^) denotes significance (n = 3, p < 0.05) in comparison to all groups in the same time point

Transfection efficiency was measured in terms of Relative Units of Fluorescence (RUF) as a function of cell concentration. Fluorescence measurements indicate that the highest efficiency was obtained with N/P ratios between 1.7 and 3.5 for HEK cells (Fig. 3B). Those results were confirmed by fluorescent microscopy, where a correlation between fluorescence measurements and fluorescent images was found. Fig. 3C correspond to a N/P ratio of 1.7, while Fig. 3D and Fig. 3E correspond to 2.6 a 3.5 N/P ratio respectively.

### WJ-MSCs transfection

After the evaluation of transfection efficiency in HEK cells, WJ-MSCs were also analyzed to establish the most suitable transfection conditions. Due to the heterogeneity of cells from human donors, it was necessary to calculate the specific growth rate of WJ-MSCs. The effect of the seeding density (C0) on the growth kinetics of WJ-MSCs is presented in Fig. 4A. When cell culture starts with a seeding concentration of 2000 cells/cm^2^ (approximately 5% confluence) an exponential growth is observed with a doubling time of 40.5 hours; however, if the initial concentration is 9000 cells/cm^2^, cell growth follows a straight line, which means that the duplication time varies from point to point and three different duplication times are obtained. At the beginning of the culture, the duplication time was 19 hours, then 44 hours and with time the crop slows down to duplication times of 85 hours. With the growth kinetics information, a seeding density of 9000 cells/cm^2^ was established for all the transfection assays, which were carried out 24 hours after cell seeding. Transfection was also evaluated 24 hours after the treatment with PEI complexes. Given that the duplication time between 24 and 48 hours is 19 hours, it is possible to infer that when the transfection is evaluated, the cells have duplicated at least once, which facilitates the entry of the complexes into the cell. Fig. 4B summarize cytometry results. N/P ratios between 3.5 and 7.0 showed transfection efficiencies between 10 and 15%, without significative differences among them. A decrease in PEI concentration reduced both efficiency and cytotoxicity, while an increase in PEI concentration also increased toxicity and an improvement in the transfection efficiency was not observed. Flow cytometry results were contrasted with fluorescence images of transfected cells. Fig. 4C corresponds to transfection with a complex formed with a N/P ratio of 1.7, while Fig. 4D and Fig. 4E correspond to 3.5 a 5.2 N/P ratio, respectively.

**Fig. 4.**
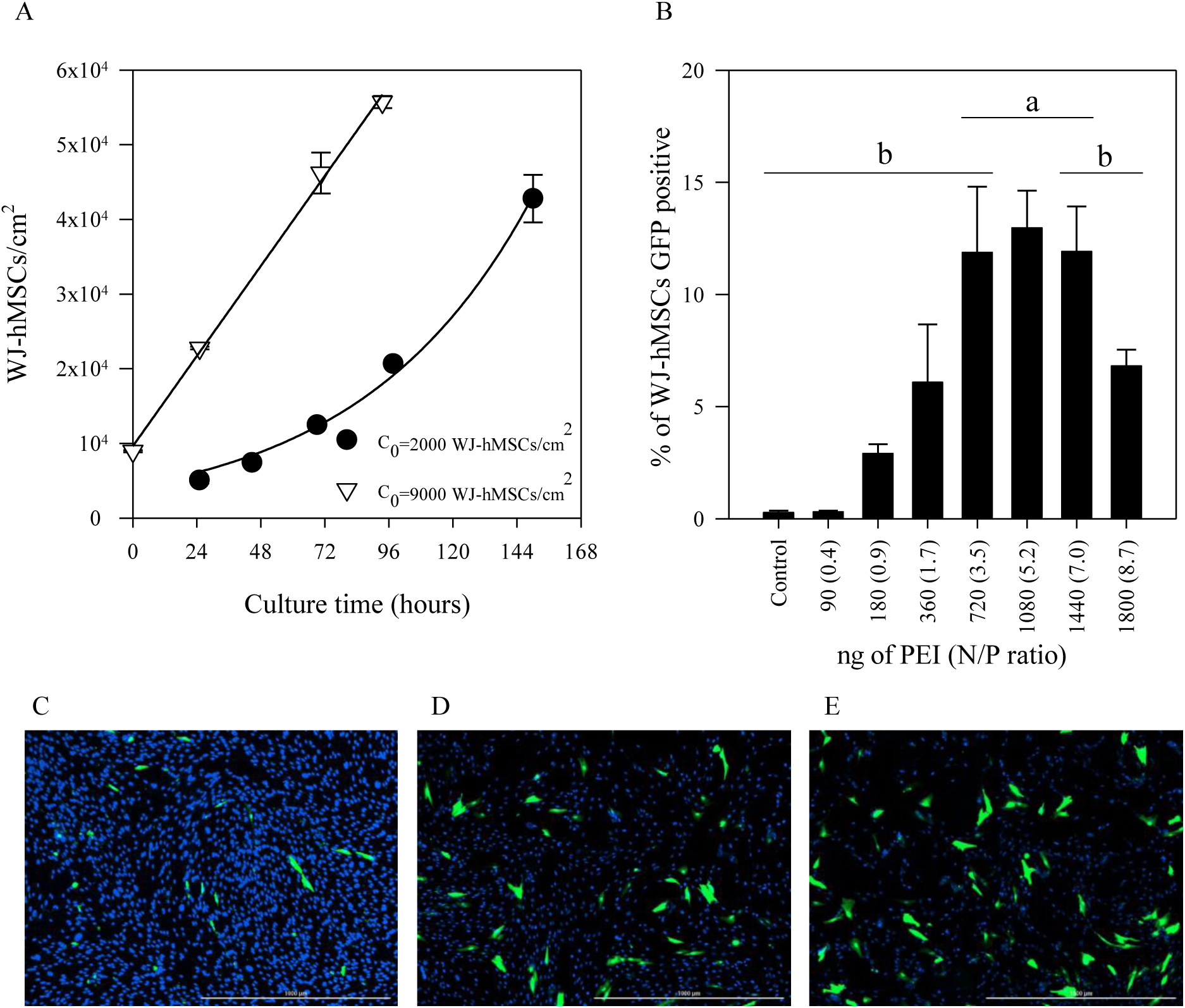
Effect of PEI concentration in WJ-MSCs transfection. (A) Effect of seeding density(C0) in growth kinetics of WJ-MSCs. (B) Relative expression levels of green fluorescent protein (GFP) by flow cytometry. (C, D, and E) Fluorescent microscope images of the GFP expressing WJ-MSCs 48 h post transfection. N/P ratios of 1.7, 3.5 and 5.2 respectively. Scale bar, 1000 um.

### Effect of PEI transfection in functional properties of WJ-MSCs

Following the international criteria for defining multipotent mesenchymal stromal cells established by the Society for Cell and Gene Therapy (ISCT), we evaluated the impact of PEI transfection on WJ-MSCs with PEI with four criteria: adherence to plastic in standard culture conditions, specific surface antigen (Ag) expression, multipotent differentiation potential (osteoblasts and adipocytes) demonstrated by staining of in vitro cell culture [24] and immunomodulatory properties. The expression of specific markers in WJ-MSCs transfected with PEI at N/P ratio of 3.5 was evaluated by flow cytometry. As observed in Fig. 5A the hematopoietic markers CD45, CD34 and HLA-DR were negative (≤2%) in both WJ-MScs transfected (T) and non-transfected (NT) WJ-MSCs, while markers for multipotent mesenchymal stromal cells (HLA-AB, CD105, CD73 and CD90) were positive (≥95%).

**Fig. 5.**
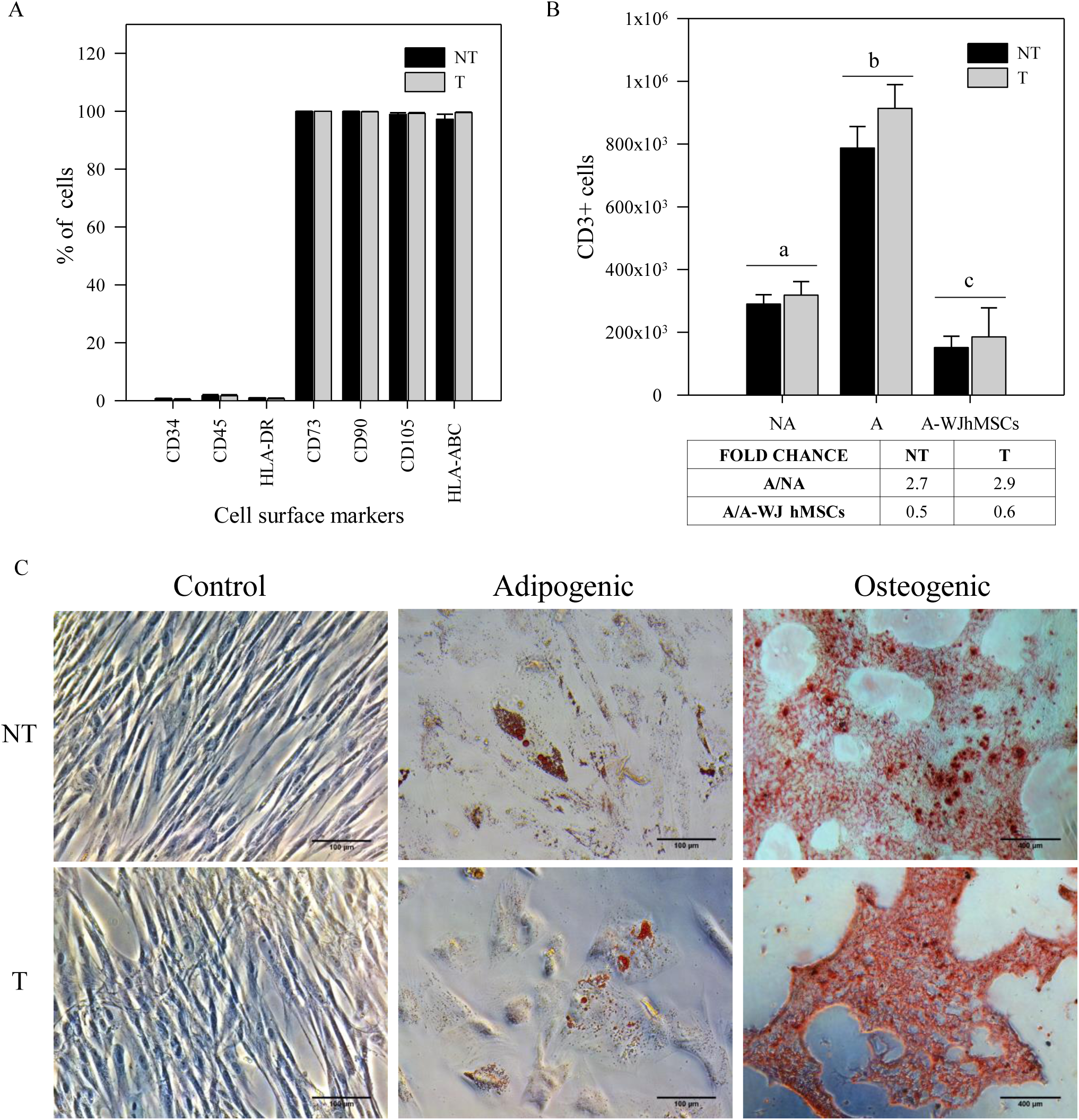
Transfection with PEI does not affect functional properties of WJ-MSCs. (A) Percentage of WJ-MSCs that express characteristic surface cell markers of mesenchymal stromal cells (B) Immunogenic activity of WJ-MSCs over PBMCs. WJ-MSCs were cocultured with PBMCs activated (A) with CD2, CD3 and CD28. Non-activated (NA) PBMCs were used as control. (C) Adipogenic (scale bar: 100 μm) and osteogenic (scale bar: 400 μm) differentiation of WJ-MSCs transfected (T) with PEI at N/P ratio of 3.5 (720ng PEI).

To evaluate the immunomodulatory capacity WJ-MSCs were co-cultured with peripheral blood mononuclear cells (PBMCs) activated (A) with CD2, CD3 and CD28. Fold change was expressed as the ratio between PBMCs activated (A) and PBMCs activated and co-cultured with mesenchymal stromal cells (A/A-WJ hMSCs). Non-activated PBMCs were used as control (NA). In Fig. 5B are presented the number of cells CD3+ (specific marker for lymphocytes) for each assay. We found that WJ-MSCs in co-culture with activated PBMCs were able to reduce CD3+ cell proliferation between 50 to 60% relative to the control in both transfected and non-transfected cells.

A multipotent differentiation potential was evaluated in T and NT WJ-hMSCs. Cells were differentiated in two different lineages: adipogenic and osteogenic. For adipogenic differentiation we observed formation of lipid vacuoles by staining with Oil Red in both experimental groups (Fig. 5C), indicating preservation of adipogenic potential upon PEI transfection. Similarly, when osteogenic lineage differentiation was evaluated, we observed formation of calcium deposits (as analyzed by alizarin red-S) in both T and NT WJ-MSCs. Together, we could confirm that MSC subjected to PEI transfection did not lose differentiation potential and immune modulatory effects, suggesting PEI-based gene transfer as safe method for MSC gene modification.

## Discussion

The overall aim of this study was to evaluate the efficacy of polyethylenimine (PEI) as a non-viral gene delivery system that can be optimized for gene therapy in human mesenchymal stromal cells from Wharton’s Jelly [25, 26]. Since both, the plasmatic membrane of the cell and plasmids have negative charge, it is necessary to reduce the repulsion between both surfaces in order to facilitate the introduction of the DNA. One alternative is using a polymer that is capable to condense DNA, in order to protect it against degradation and, additionally, to serve as a vehicle to cross the cell membrane.

PEI binds DNA to form complexes; this interaction was evaluated thought electrophoretic motility. The electrophoresis assay showed that PEI begins complexing pGFP when the N/P ratio is greater than 0.9; however, measurements of particle size and zeta potential were only achieved with N/P ratios greater than 1.7. This suggests that N/P ratios lower than 1.7 are not enough to form stable particles. Then we evaluated if an increase in PEI concentration affected particle size, zeta potential and efficiency of transfection. We found that for both HEK and WJ-MSCs there is a N/P ratio at which the highest transfection is reached; however, the ratio and the efficiency vary from one cell type to another. In HEK cell line, the highest transfection (60%) was obtained at N/P equal to 2.6 while in WJ-MSCs was required a N/P ratio of 3.5 to reach 13% of transfected cells.

When particle size and zeta potential were analyzed, it was found that both physical properties of the complexes remained constant at N/P ratios from 1.7 to 8.7, which suggests that the variation in transfection efficiency for each of the cell types is not associated with size and charge of the particles but may be associated with free PEI that does not complex with DNA. According to Yue (2011), the excess of PEI in the solution improves the internalization of the complexes and can contribute to the subsequent intracellular traffic, however, Hanklikova et al (2011) have found that these free groups are the ones that contribute the most to the toxicity during transfection [27, 28]. When the toxicity was evaluated, it was found that for the highest efficiency of transfection in HEK (N/P = 2.6), the percentage of viability was greater than 70%, while in the WJ-MSCs it was founded that PEI reduces cell viability in 40% when its concentration is equal or superior to 720 ng (N/P=3.5) in 100 µl (7.2 µg/ml) for 4.5×10^4^ WJ-MSCs. At this value, transfection efficiency was 13±2% approximately and remained constant with the increase of PEI concentration up to a N/P ratio of 7.0, however at this value, cytotoxicity grew up considerably until 70%.

Those transfection values in WJ-MSCs are comparable with values obtained by other authors. For example, the highest efficiency in hMSCs from bone marrow founded by Wang et al (2011), with cell viability near 60%, was 25%, achieved at N/P ratio 2 and 6 μg DNA/cm^2^ [21], however they obtained this efficiency using 12000 ng in comparation with our experiment, when final plasmid concentration per well was 400 ng (30 times less).

Since between N/P ratios of 3.5 and 7.0 the same transfection efficiencies were achieved for WJ-MSCs but toxicity increased proportionality with PEI, a N/P ratio of 3.5 was selected to evaluate characteristic properties of the WJ-MSCs in order to establish if transfection with PEI affects its immunophenotype, influence its immunomodulatory properties and its differentiation capacity. According to the ISCT [15, 16], WJ derived cells can be classified under the name of stromal cells, if: 1) plastic-adherence is maintained in standard culture conditions, 2) cells express CD105, CD73 and CD90, and lack expression of CD45, CD34, CD14 or CD11b, CD79a or CD19 and HLA-DR surface molecules and 3) cells are able to differentiate to osteoblasts, adipocytes and chondroblasts *in vitro*. Our results show that the transfection of the WJ-MSCs with the PEI-pGFP complex does not affect these functional characteristics of stromal cells; that means, PEI can be used as transfection agent due to WJ-MSCs retain its stemness. Thus, in future works, GFP can be replaced with other DNA sequences in order to favor differentiation towards a specific lineage, for example SOX5, 7 or 9 in chondrogenesis process [4] or BMP-2 and runt-related transcription factor 2 (RUNX2) in osteogenesis [29]. Additionally, tissues are tridimensional structures, therefore transfection process could take place in a biocompatible porous scaffold, where cells can growth and proliferate and then, they may encounter the PEI-DNA complex into the scaffold and transfection process will be controlled by growing cells rate [30].

## Conclusions

In this study, we evaluated the potential use of PEI as transfection agent of WJ-MSCs. We prepared complexes with PEI and pGFP at different concentrations, finding an optimal value of 720 ng of PEI (N/P ratio of 3.5), which was used for the immunomodulation and differentiation assays. Our results showed that WJ-MSCs transfected with PEI retain their morphology, plastic adherence, immunophenotype, immunomodulatory function and multi-lineage differentiation potential and therefore, PEI can be used as a transfection system in applications for therapeutic purposes. The standardization of the transfection conditions of WJ-MSCs with PEI has great potential for tissue engineering applications, due to the differentiation capacity of WJ-MSCs, in future studies it will be possible to change GFP for proteins that favor differentiation osteogenic, adipogenic or chondrogenic depending on the desired application.

## Abbreviations

Ag: Specific surface antigen
DAPI: 4′,6-diamidino-2-phenylindole
DLS: Dynamic Light Scattering
DMEM: Dulbecco’s Modified Eagles Medium
DMSO: Dimethyl sulfoxide
DW: Distilled water
FBS: Fetal Bovine Serum
GFP: Green fluorescent protein
HCl: chloride acid
HLA: Human leukocyte antigen
HLA-AB: HLA class I molecules are present in all nucleated cells in human. This region includes HLA-A, -B, -C, -E, -F, -G, -H, -J, and HLA-X loci. HLA-A, HLA-B, and HLA-C loci are polymorphic and functional classical class I loci
HLA-DR: HLA class II molecules are located on B lymphocytes, macrophages, dendritic cells, endothelial cells, and active T cells. They consist of six different loci (HLA-DM, DN, DO, DP, DQ, and DR).
hMSCs: Human mesenchymal stromal cells
ISCT: International Society for Cell and Gene Therapy
LB: Luria Bertani
MSCs: Mesenchymal stem cells
N/P ratio: N correspond to the nitrogen in PEI and P refers to phosphate content in DNA.
NADH: Nicotinamide adenine dinucleotide exists in two forms: an oxidized and reduced form, abbreviated as NAD+ and NADH
NADPH: NADPH is the reduced form of NADP+ (Nicotinamide adenine dinucleotide phosphate)
OD600nm: Optic density at 600 nm
PBMCs (A): Peripheral blood mononuclear cells activated
PBMCs (A/A-WJ hMSCs): Peripheral blood mononuclear cells activated and co-cultured with mesenchymal stromal cells
PBMCs (NA): Peripheral blood mononuclear cells and without activation
PBMCs: Peripheral blood mononuclear cells (PBMCs)
PBS: Phosphate Buffered Solution
PEI: Polyethylenimine
pGFP: Plasmid codifying for green fluorescent protein
RFU: Relative Fluorescence Units
TAE: Tris-acetate-EDTA
TGF-β: Transforming Growth Factor-β
UC-hMSCs: Umbilical cord human mesenchymal stromal cells
WJ-MSCs: Wharton’s Jelly-derived mesenchymal stromal cells
WJ-MSCs (NT): Wharton’s Jelly-derived mesenchymal stromal cells non-transfected
WJ-MSCs (T): Wharton’s Jelly-derived mesenchymal stromal cells transfected with polyethylenimine

## Declarations

## Acknowledgements

AR thanks Margareth Patiño and Leslie Sanchez for their help in *E. coli* DH5-α transformation and plasmid extraction, respectively.

## Funding

This work was supported by Universidad Nacional de Colombia under Grant 203010026990 and Departamento Administrativo de Ciencia, Tecnología e Innovación (COLCIENCIAS) through the Doctoral Scholarship Program 567-2012.

## Availability of data and material

All data generated or analyzed during this study are included in this published article.

## Authors’ contributions

AR conceived, design and performed the experiments and analyzed the data. ER performed the experiments of electrophoresis, zeta potential and size of complexes. CR and KB performed the experiments of immunophenotype, immunomodulation properties and multipotential capacity of WJ-MSCs. BC and GS participated in data analysis and interpretation. RG conceived and designed the experiments and was responsible for critical review of the manuscript. All authors were involved in revising the manuscript, and the final manuscript was read and approved by all authors.

## Ethics approval and consent to participate

Written informed consent for the use of the umbilical cord was provided from each woman.

## Consent for publication

Not applicable

## Competing interests

The authors declare that they have no competing interests

## Notes

### Competing Interest Statement

The authors have declared no competing interest.

